# MOCCASIN: A method for correcting for known and unknown confounders in RNA splicing analysis

**DOI:** 10.1101/2020.06.16.154674

**Authors:** Barry Slaff, Caleb M Radens, Paul Jewell, Anupama Jha, Nicholas F Lahens, Gregory R Grant, Andrei Thomas-Tikhonenko, Kristen W. Lynch, Yoseph Barash

## Abstract

While the effects of confounders on gene expression analysis have been extensively studied there is a lack of equivalent analysis and tools for RNA splicing analysis. Here we assess the effect of confounders in two large public RNA-Seq datasets (TARGET, ENCODE), develop a new method, MOCCASIN, to correct the effect of both known and unknown confounders on RNA splicing quantification, and demonstrate MOCCASIN’s effectiveness on both synthetic and real data.

## Introduction

RNA-Seq is an experimental technique that quantifies the relative abundance of RNA molecules in a sample through sequencing of short RNA fragments. By mapping those sequences to a transcriptome or an annotated genome, researchers can quantify expression levels and alternative splicing of genes. RNA-Seq is commonly used in a variety of analysis tasks like the identification of differentially expressed genes or isoforms between two or more conditions; the identification of genes and/or samples that cluster together by gene expression or RNA splicing variations; and quantitative trait loci analysis to identify genetic variants associated with changes in expression (eQTL) or splicing (sQTL). The results of such analyses can be greatly affected by unwanted factors such as sequencing lane (Lin et al. 2014; Bullard et al. 2010) or processing batch (Busby et al. 2011; Peixoto et al. 2015). These factors are either already known prior to starting the analysis task (e.g. sequencing lane), or unknown (e.g. difference in mouse diet which was never recorded). Generally, such confounders (also termed nuisance variables) affect the variability of the data and, depending on the relationship to the biological signal of interest, can lead to inflated rates of false positives and false negatives.

Fortunately, for gene expression analysis, there are well-established methods to remove or account for both *known* and *unknown* confounding factors, such as limma (Smyth and Speed 2003), ComBat (Johnson, Li, and Rabinovic 2007), RUV (Risso et al. 2014; Gagnon-Bartsch and Speed 2012), and surrogate variable analysis (svaseq) (Leek and Storey 2007; Leek 2014). All of these tools have been applied in many studies and are highly cited.

In contrast, there is a clear lack of equivalent tools for modelling *known* and *unknown* confounding factors in alternative splicing analysis. We suspect this scarcity of tools reflects a general lack of awareness of the effect of confounders on splicing analysis. Specifically, we were not able to find any previous work that quantitatively assessed the effect of confounders on splicing analysis and compared it to the effect on gene expression.

One challenge with correcting for confounders in RNA splicing analysis is that tools designed for gene expression analysis are not well suited for this task. Local splicing variations (LSV) are typically quantified from junction reads, a subset of exon-mapped reads spanning across introns. When junction reads from a reference exon map to two or more alternative splice sites up/downstream, the percent splice inclusion (PSI) of each of these splice sites is quantified as the ratio of junction reads mapping to that splice site over the total number of junctions reads mapped to all splice sites at that locus. Similarly, between samples or groups of samples, the delta PSI (dPSI) quantifies the extent of changes in PSI, or differential splicing. Consequently, alternative splicing (AS) quantification is distinct from that of gene expression: rather than real values or log fold change, PSI values are in the range of 0 to 1, or -1 to +1 for dPSI. LSV quantification also suffers from different biases and challenges than those of gene expression and therefore algorithms tailored for their quantification give better PSI and dPSI estimates than algorithms that infer full isoforms expression levels (c.f (Vaquero-Garcia et al. 2016a), SUPPA (Alamancos et al. 2015)).

Most previous work involving splicing analysis and confounding factors utilized pipelines originally built for QTL analysis. For example, in (Raj et al. 2018) the authors quantified variations in intron cluster usage using LeafCutter (Li et al. 2018) and then applied fastQTL (Ongen et al. 2016) to the data matrix. This analysis implicitly assumes gaussian distributions, standardizes the data, and then corrects for confounders such as age or gender. However, not only do PSI values not follow a Gaussian distribution, but the corrected values are commonly negative or greater than one, losing their interpretation as splicing fractions. Furthermore, these pipelines use only point estimates for PSI, losing read coverage information which is crucial for controlling false positives in differential splicing analysis (Zhao et al. 2013). Finally, while some software allow users to specify confounders for differential splicing analysis (Zhao et al. 2013; Anders, Reyes, and Huber 2012; Reyes et al. 2013; Li et al. 2018), we are not aware of any tool that is able to correct for both known and unknown confounders, outputting corrected junction spanning reads for both supervised (e.g. differential splicing) and unsupervised (PSI clustering) splicing analysis tasks.

In order to address the lack of tools for correcting confounders in RNA-Seq based splicing analysis we introduce MOCCASIN (Modeling Confounding Factors Affecting Splicing Quantification). We evaluate MOCCASIN’s performance on both synthetic and real datasets, demonstrating that it works effectively to eliminate false positives and recover true biological signals diminished due to confounders.

## Results

First, we assessed the effect of confounders on both expression and splicing analysis in two large and highly used datasets each with hundreds of samples. As shown in Fig1 we found the effect of confounders on splicing analysis is at least as large as the effect on expression analysis. Specifically, in the TARGET’s acute lymphoid leukemia dataset and ENCODE’s shRNA knockdown (KD) dataset the batch labels were associated with 15.9% and 46% respectively of the total splicing variations compared to 6.7% and 39.8% for expression. Indeed, the authors of (Nostrand et al. 2018) noted such batch effects for ENCODE, but this effect was not quantified and we are unaware of a previous publication reporting batch effect associated with the sequencing machine for the TARGET dataset. Importantly, batch effects are not restricted to such large consortium data, with similar results observed for mouse tissue samples previously used to study expression batch effects by (Peixoto et al. 2015) (See Supplemental Figure 1).

**Figure 1:**
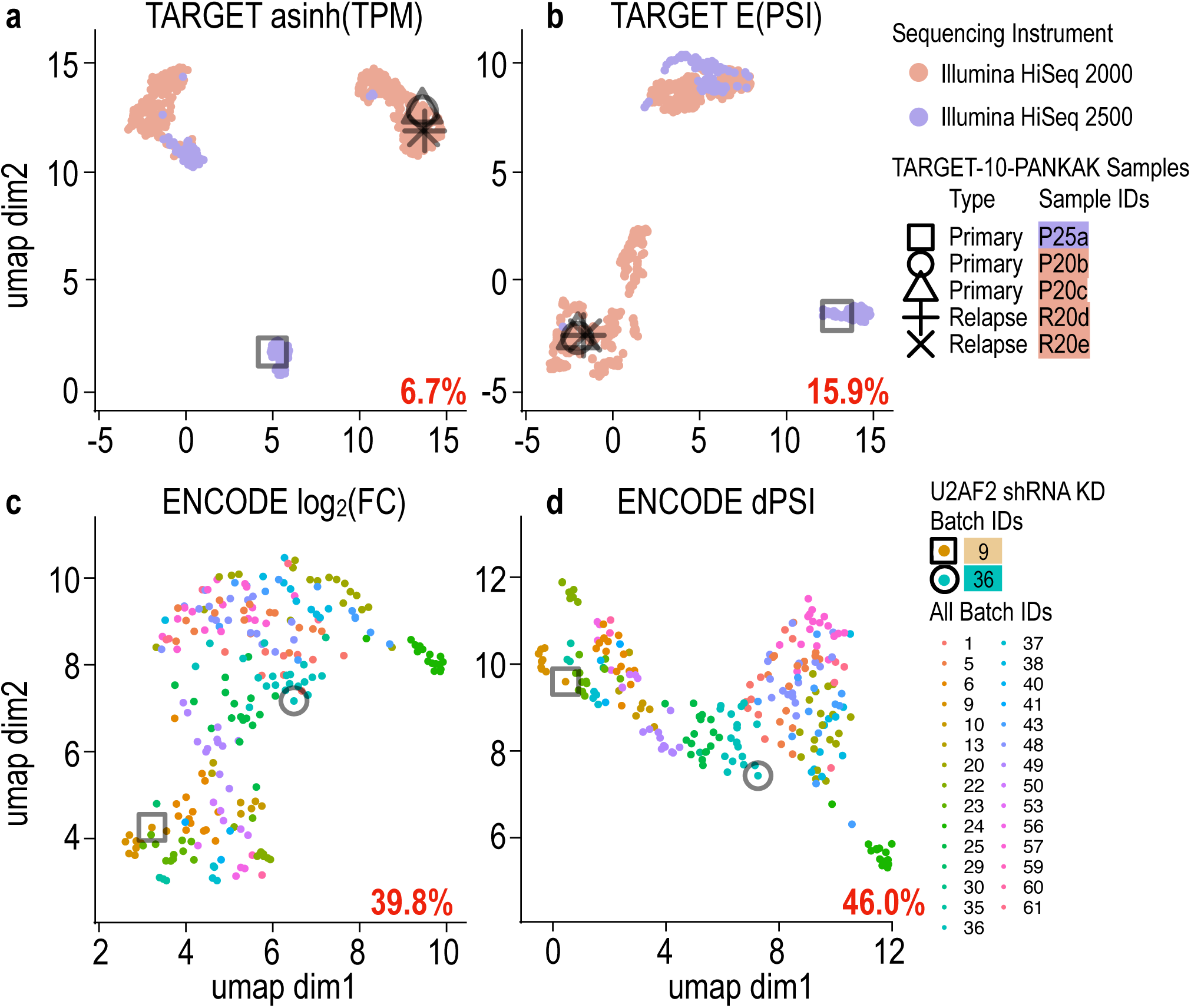
Batch effects impact both gene expression and splicing analysis. Uniform manifold approximation and projection (umap) of gene expression analyses (**a, c**) and splicing analysis (**b, d**) for TARGET (top) and ENCODE (bottom). Colors indicate batch identity. Numbers in red represent total variation associated with batch in each dataset. Shapes mark samples from the same patient (TARGET, patient TARGET-10-PANKAK) or experiment type (ENCODE, U2AF2 KD) which cluster by batch.

In order to address the need for correcting confounding factors in splicing analysis we developed MOCCASIN. MOCCASIN operates by adjusting the number of junction spanning reads jointly for each LSV. Briefly, for each junction *m* in sample *k*, design matrices of known confounders *N*, unknown confounders *U*, and variables-of-interest *V*, we assume:

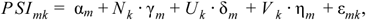

Such that ordinary least squares (OLS) regression can be completed independently for each junction. With 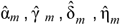 denoting the estimated coefficients we have:

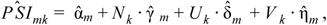

Such that *r*_*mk*_ := *PSI*_*mk*_ - *P ŜI*_*mk*_ is the residual, 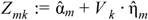 is PSI predicted from the *partial model* which excludes confounding factors, and *A*_*mk*_ := *r*_*mk*_ + *Z*_*mk*_ is the adjusted PSI. In the final adjustment step, negative PSI are clipped to 0, PSI are re-normalized for each LSV so that the sum over junctions is 1 for each sample, and adjusted counts for each junction are computed by multiplying the adjusted PSI into the initial total coverage for that LSV and sample.

MOCCASIN is designed to work with MAJIQ’s LSV quantifications. Specifically, MAJIQ produces bootstrap samples for each junction coverage in dedicated *.majiq files, which serve as MOCCASIN’s input. It then performs the above adjustment per bootstrap sample and outputs the adjusted *.majiq files. The adjusted *.majiq files can then be fed into MAJIQ for detecting differential splicing between conditions of interest or into unsupervised clustering algorithms as described below.

We first applied MOCCASIN to synthetic data, generated by BEERS (Grant et al. 2011), to simulate 16 samples of RNA-Seq produced by (Zhang et al. 2014) from mouse tissues (Aorta and Cerebellum). Running MAJIQ on these data produced the results labeled as ground truth with respect to the biological signal of interest (tissue difference) as well as any detected difference between two randomly chosen subsets of the samples labeled as batches A and B (batch difference). We then injected an artificial batch signal into *G* percent of randomly chosen genes as follows: First, we reduced the TPM of the most highly expressed isoform of the gene by *C* percent. Next, we randomly selected another of the gene’s isoforms and increased its expression by the same amount, thus maintaining the gene’s overall original TPM. Fig. 2a shows the batch effect on the number of detected differentially spliced LSV (y-axis) as a function of the difference’s significance (x-axis, Wilcoxon *p*-value, see Methods). Here, with *G*=20% and *C*=60%, we see the batch injected data (blue) causes inflation of both batch associated differences (Fig. 2a) as well as false positives for the tissue signal (Fig. 2b). Applying MOCCASIN (orange) effectively controls both the batch signal and the false positives for the biological signal with only a moderate loss of true biological signal (false negative) due to over correction at the moderate p-value levels. To better quantify these effects across a range of settings we applied MAJIQ’s Bayesian dPSI model to report all LSVs for which the posterior probability of a PSI change of above 20% is at least 95% (P(|dPSI| >= 20%) >= 95%). This threshold is commonly used to detect splicing changes that are considered biologically significant, of high confidence, and can be validated using orthogonal approaches such as RT-PCR. Fig2c-d show that small perturbations of 2% or even 10% have little effect on the false positive rate (FPR) with respect to the original batch signal (Fig. 2c) and the false discovery rate (FDR) with respect to the tissue signal (Fig. 2d). These results are to be expected given the dPSI thresholds set for MAJIQ, reflecting its robustness to small perturbations below the specified detection threshold. However, the picture changes dramatically when the injected signal reaches *C*=60%, with both FPR for batch signal and FDR for tissue signal climbing proportionally as *G* increases from 2% to 5% and 20% (blue). However, applying MOCCASIN (orange) effectively controls for the batch effect and perhaps most importantly maintains an empirical FDR of 6-8%, close to the Bayesian estimate of 95% confidence. We also found that using known batch covariates or treating those as unknown and letting MOCCASIN correct them performed similarly (grey), and this result was not sensitive to the number of unknown confounders used (compare U1, U2, U3 in Supplementary Figure 2). Finally, the overall effect on downstream unsupervised analysis tasks is demonstrated in Fig2e where the original data clusters by tissues (left), by batch when the batch signal is injected (middle), and again by tissue after MOCCASIN is applied (right).

**Figure 2:**
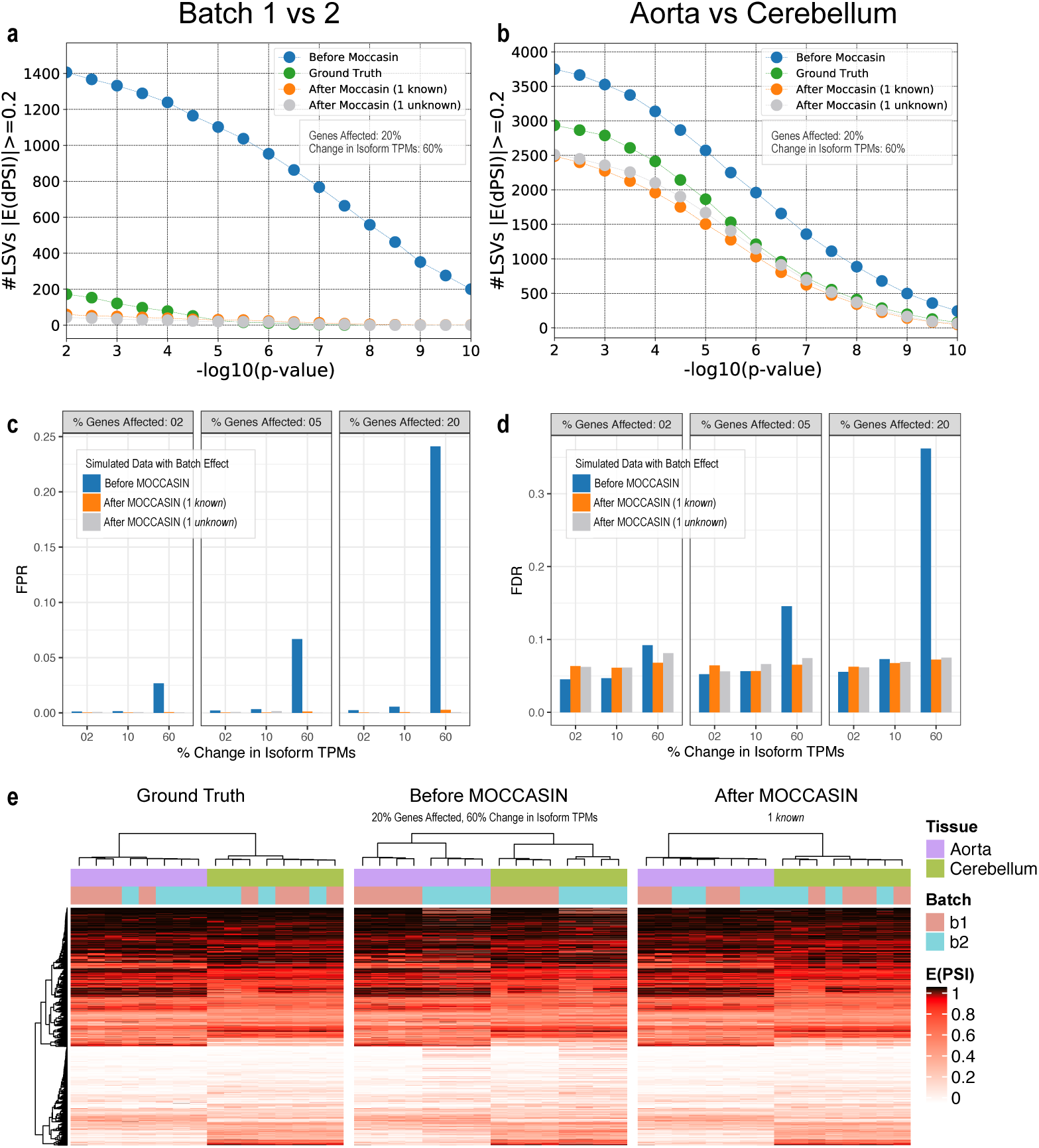
Removal of batch effects from simulated RNA-Seq data with MOCCASIN. RNA-Seq samples from Aorta and Cerebellum were simulated using BEERS while injecting *G*% of the genes in half the samples with a batch effect of *C*% expression change of the main isoform (see main text). **a**,**b)** The number of LSVs (Y-axis) detected as differential (|E(dPSI)|>0.2) for the batch signal (a, left) and the tissue signal (b, right) across a range of increasingly significant p-values (X-axis, Wilcoxon rank sum test, -log_10_ scale). The green (with batch signal injection *G*=20%, *C*=60%) and blue (no batch signal injection) points are from the simulated data without MOCCASIN correction and serve as a reference. Orange and grey represent respectively the results after MOCCASIN correction when the batches are known or unknown. **c**,**d)** Assessing false positive rate (FPR) for the batch signal (c, left) and false discovery rate (FDR) for the tissue signal (d, right) for a range of *G* values (2,5,20%) and *C* effect size (2, 10, 60%). Here positive events where considered as those changing by at least 20% with high confidence by MAJIQ (P(|dPSI|>20%)>95%). Under these definitions small effect size (*C*=2,10%) represent perturbations which are not expected to affect the positive event set much. **e)** Heatmaps of E(PSI) from simulated data without batch effect (ground truth, left), with simulated batch effect (G=20%, C=60%) without correction (middle), and after applying MOCCASIN with 1 known confounding factor (right). Each column is a sample, and each row is an LSV. The colored bars above the samples denote the sample’s tissue (aorta purple, cerebellum green) and (batch 1 red, batch 2 blue).

The application of MOCCASIN to the TARGET and ENCODE datasets described above are shown in Fig3. In both datasets we find that after applying MOCCASIN the amount of variance explained by batch drops significantly, from 15.9% to 0.4% in TARGET and from 46% to 4.3% in ENCODE. However, detecting differences between samples that represent true biological signals is generally maintained, or even improved by as much as 12% when applying MOCCASIN. For example, ENCODE targeted a key spliceosomal factor U2AF2 for knockdown (KD) with shRNA in two distinct experiment batches: 9 and 36. In batches 9 and 36, we identified 2595 and 1897 U2AF2 KD-induced changes in splicing (|dPSI|>=0.1, p-value < 0.05), respectively, of which only 773 were identified in both batches (Fig. c, left). After applying MOCCASIN (Fig. c, right) to model and remove batch-associated variation, the overlap between batch 9 and 36 increased from 773 to 1048, the overall Pearson correlation between batches increased from 0.71 to 0.79, and the total number of detected U2AF2 KD-induced changes increased by 12.2% and 8.1% for batches 9 and 36, respectively. Comparing a within-batch analysis (same sequencer) to a mixed-batch (multiple sequencers) analysis also showed that TARGET data corrected by MOCCASIN correlated better (Fig. 3d, top: Pearson correlations increased from 0.87 to 0.95) and overlapped more (Fig. 3d, bottom: overlap increased by 10.1%) when detecting patient-specific primary ALL diagnosis versus relapse-associated splicing differences (P(|E(dPSI)|>=0.1)>=0.95).

**Figure 3:**
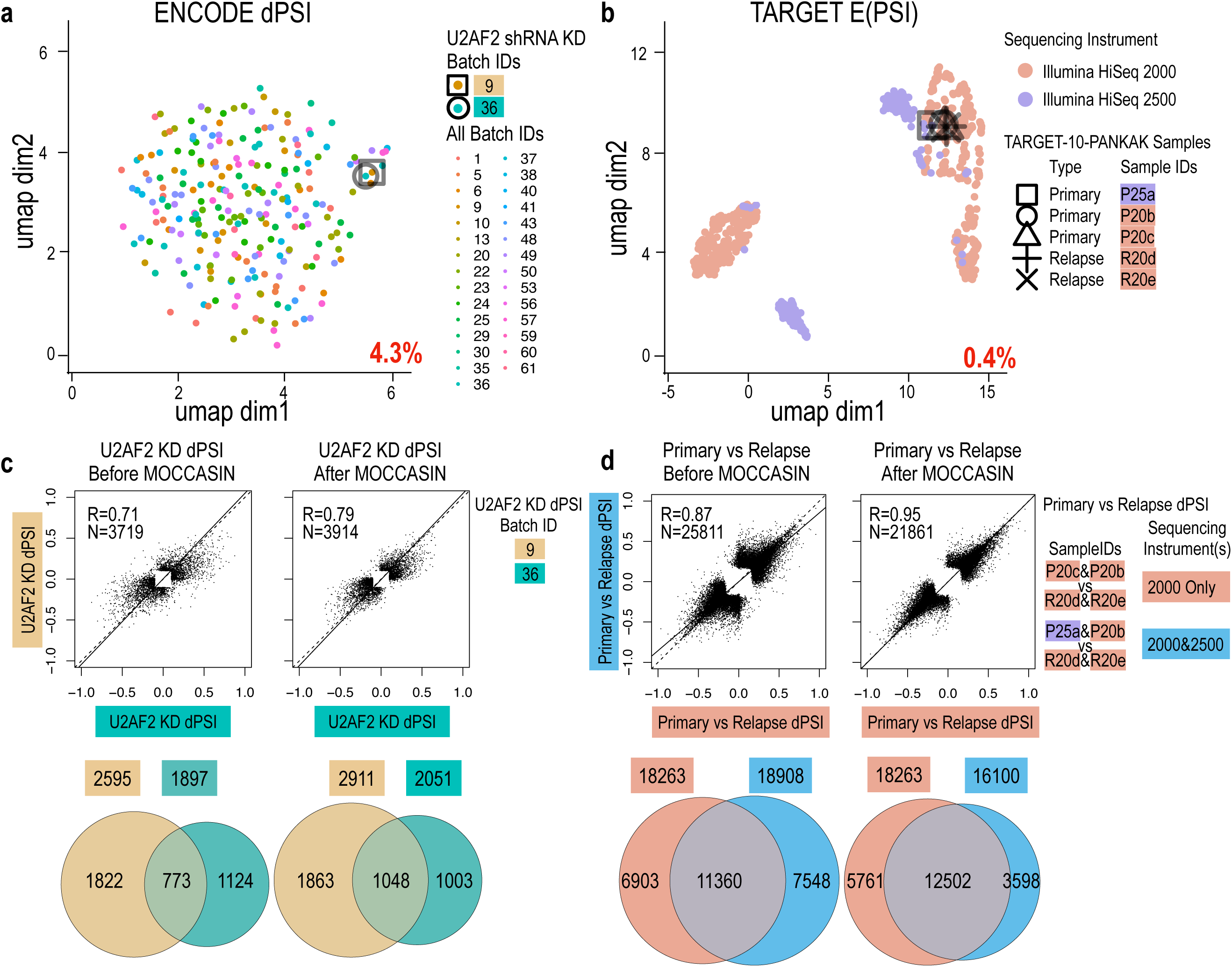
Batch correction of TARGET and ENCODE datasets. **a**-**b**, umap plots as in Figure 1 but after applying MOCCASIN. U2AF2 knockdowns (KD) now cluster together (square and circle in **a**) and TARGET technical replicates P25a-c cluster together (square, circle, and triangle in **b**). The percent of total variance due to sequencer (**a**) or experimental batch (**b**) across each dataset is indicated in red in the bottom-right corner. **c)** Correlation of significant splicing changes upon U2AF2 KD (dPSI > 20%) between batches increases from R=0.7 (top left) to R=0.79 after MOCCASIN (top right). Similarly, the number of significantly changing events (P(|dPSI|>20%)>95%) in each batch and the overlap of these events between batches increases after MOCCASIN correction (c, compare bottom left and right). (**d**) Correlation of significant splicing changes between primary and relapse samples from the same patient (TARGET-10-PANKAK) increases from R=0.87 (top left) to R=0.95 (top right) after correcting for the sequencing platform. Note that since only the HiSeq2500 samples are corrected by MOCCASIN, only the set that includes those (cyan) is affected. Accordingly, the total number of significantly changing events (P(|dPSI|>20%)>95%) drops for the cyan set from 18908 (bottom left) to 16100 (bottom right) but the overlap with the other set increases (11360 to 12502).

## Discussion

While the issue of confounding factors’ effect on gene expression analysis has received much attention, the equivalent effect of confounders on RNA splicing analysis has not been well studied. Here we use two large public datasets to show the magnitude of confounders’ effect on RNA splicing quantification is at least as big as that observed for gene expression analysis. We develop MOCCASIN, the first dedicated tool to correct for both known and unknown confounders in RNA splicing analysis and demonstrate its effectiveness on both synthetic and real data. We have also made all code and data available. Specifically, in addition to correcting ENCODE HepG2 data (Fig3), we also corrected the K562 data (Supplementary Figure 3). The corrected HepG2 and K562 ENCODE dataset should serve as a highly valuable resource as it offers, for the first time, quantification of both complex and de-novo splicing variations which were not available before, as well as MOCCASIN batch corrections. Similarly, the synthetic data generated for this study (see supplementary methods) can serve as a benchmark for future tool development.

While we focused here on large public datasets it is important to note similar observations can be made in much smaller datasets. For example, the data from (Leek 2014) which were used in the development of RUV for expression correction, exhibited strong batch effects in splicing analysis as well (see Supplementary Figure 1). Another important point to make is with respect to scalability. Nowadays datasets can easily involve hundreds or even thousands of samples, requiring efficient algorithms. This problem is further compounded by the fact that the number of junctions in the human transcriptome is typically an order of magnitude larger than the number of genes. Consequently, we chose to utilize a relatively simple model which appears to perform well while still scaling to large datasets as those used here. However, more elaborate models could be introduced in the future to better model specific effects. These may include non linear models, mixed effects models, and models which share information across the corrected LSVs.

We hope that the combined code, data, and analysis we provided here will serve as a valuable resource for the research community and will shed light on the need to control for confounders in RNA splicing analysis.

## Methods

### Data Availability

The TARGET results published here are in whole or part based upon data generated by the Therapeutically Applicable Research to Generate Effective Treatments (https://ocg.cancer.gov/programs/target) initiative, phs000218. The TARGET data used for this analysis were accessed under Project #10088: Alternative splicing in pediatric cancers (request 41466-5). The SRA toolkit was used to download sra files for the TARGET dataset and simulated data was based off of data from (Zhang et al. 2014) (GEO accession GSE54652). The ENCODE project fastqs were downloaded from www.encodeproject.org.

### Introducing batch effects by perturbing TPMs

Batch effects were introduced to the transcript level TPMs of four aorta and four cerebellum samples. The first four of the aorta samples and the first four of the cerebellum samples were defined as “batch 1.” The last four of the aorta samples and the last four of the cerebellum samples were defined as “batch 2.” Batch effect perturbations were only ever introduced to the batch 2 samples and batch effects were restricted to genes with at least two protein coding transcripts and at least one transcript with ≥ 10 reads per kilobase of transcript length in every sample. The procedure to introduce batch effects is as follows: first, the most abundant protein coding transcript per gene was identified as the transcript having the maximum over all transcripts of the minimum reads per kilobase over all samples. This definition ensures the selected transcript is not zero in any sample. Then, for a given gene a batch effect was introduced by (1) selecting a transcript uniformly at random (excluding the most abundant transcript) and (2) reducing TPM of the most abundant transcript by a factor of ‘*C*% Change in Isoform TPM’ and correspondingly increasing the TPM of the randomly selected transcript, thus maintaining the overall TPM of the gene and not breaking the definition of TPM (sum of all TPMs in a sample is one million). The ‘*C*% Change in Isoform TPM’ factors included 2%, 10%, and 60%. In addition to introducing three different levels of percent changes in isoform TPMs, we also varied the percent of genes batch-effected. We introduced batch effects to *G*=0%, 2%, 5% or 20% of all genes with protein coding transcripts.

### Simulated data generation

The mouse cerebellum and aorta simulated data are based on the real mouse cerebellum and aorta data from (Zhang et al. 2014). We used transcript-level TPM quantifications from these data as the empirical distributions for simulating RNA-Seq reads with the Benchmarker for Evaluating the Effectiveness of RNA-Seq Software (BEERS) (Grant et al. 2011). Briefly, we used BEERS to simulate 30e6 paired end RNA-Seq reads for each input sample, with uniform coverage across the length of each expressed transcript and no intronic expression. We also introduced polymorphisms and errors into the resulting data with the following parameters: substitution frequency = 0.001, indel frequency = 0.001, and error rate = 0.005.

### MOCCASIN algorithm methods

MOCCASIN adjusts junction counts representing evidence for RNA splicing events in order to remove confounding variation. The main inputs provided by the user are a model matrix and list of confounding factors (subset of the model matrix columns). The model matrix identifies the input samples to be processed by MOCCASIN. In the simplest case, each input sample is represented by a coverage table, with each row of the table representing a splice junction and the row value corresponding to read-counts providing evidence for that junction. MOCCASIN considers the fundamental splicing unit of interest to be the local splicing variation (LSV), where each LSV refers to a collection of splice junctions with either a common source or target exon (Vaquero-Garcia et al. 2016b; Norton et al. 2018). Hence, the junctions represented in the coverage table are assumed to be grouped into LSVs, with a separate data structure identifying rows with LSVs. MOCCASIN aggregates the junction read evidence from the input samples into a table with row dimension M x K, where M is the number of common LSV junctions and K is the number of input samples.

For each LSV and sample, MOCCASIN first computes the total coverage (sum of reads over junctions in the LSV). Then the counts are converted to PSI (proportion spliced-in), i.e. proportion of counts per junction from the total. Each PSI value is in [0, 1] and PSI values over all junctions in an LSV sum to 1. An exception is the case in which an LSV has 0 coverage for some sample; in this case the “PSI” values are set to 0 for the junctions (and hence do not sum to 1).

The *M* x *K* junction PSI values together with the model matrix are the inputs to the adjustment step. Ordinary least squares regression is used to find estimators for the model terms. Specifically, for junctions indexed 1…*M* and samples indexed 1…*K*, we model

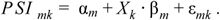

In the above,

- α_*m*_ is the intercept for junction *m*
- *X*_*k*_ is the model matrix row corresponding to the *k*’th input sample
- β_*m*_ are the coefficients for junction *m*
- ε_*mk*_ is the error term

OLS regression is completed independently for each junction to find estimators 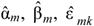 for each *k.* Thus, each OLS regression problem includes *K* observations (one per sample).

For the purposes of adjusting PSI, we partition the columns of the model matrix *X* into two groups, corresponding to confounders *C* and variables-of-interest/non-confounders *V*. In the case when the known confounders are supplemented with additional factors of unwanted variation (using for example the RUVr approach as described below), the columns of *C* are partitioned into known *N* and unknown *U* confounders. Then the expression above may be rewritten

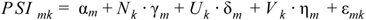

Here, γ, δ, η are the coefficients corresponding to *N, U, V* respectively.

Once fit, this constitutes a learned model for junction *m*. Denote by 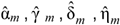 the learned coefficients:

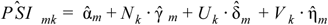

The adjustment procedure for PSI is then executed by computing:

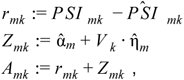

so that:

- *r*_*mk*_ is the residual
- *Z*_*mk*_ is PSI predicted from the *partial model* which excludes confounding factors
- *A*_*mk*_ is the adjusted PSI

In the final adjustment step,

1. Negative PSI are clipped to 0
2. PSI are re-normalized for each LSV so that the sum over junctions is 1 for each sample. As before, LSVs which are all-zero (after adjustment) remain 0.
3. Adjusted counts for each junction are computed by multiplying the adjusted PSI into the initial total coverage for that LSV and sample.

## Acknowledgements

This work was supported by R01 GM128096 to Y.B; U01 CA232563 to Y.B., A.T.T, and K.W.L

## Author Contributions

Conceptualization: Y.B., B.S.; Methodology: Y.B., B.S.; Software: B.S., P.J.; Investigation: Y.B., C.M.R., B.S.; Formal Analysis: Y.B., C.M.R., B.S., A.J.; Synthetic Data Generation: N.F.L, G.R.G.; Data acquisition and curation: C.M.R, A.T., Y.B.; Visualization: C.M.R., B.S. Writing, Review and Editing: Y.B., C.M.R., B.S., K.W.L., G.R.G, N.F.H, A.T; Writing, Original Draft: Y.B., C.M.R.; Supervision: Y.B., K.W.L;

## Code Availability

Moccasin code will be made available upon publication. If you would like to run moccasin on your data prior to publication please contact.

