## Supplemental Methods for "MOCCASIN: A method for correcting for known and unknown confounders in RNA splicing analysis"

1 MOCCASIN Supplementary Methods 2
1.1 Data Availability 2
1.2 RNA-Sequencing data processing steps 2
1.3 Introducing batch effects by perturbing TPMs 2
1.4 Simulated data generation 3
1.5 ENCODE dataset analysis 3
1.6 Gene expression analysis 4
1.7 Splicing analysis 4
1.8 Uniform manifold approximation and projection (UMAP) 4
1.9 Hierarchical clustering 4
1.10 Percent of total variance associated with batches 5
1.11 Scatterplots and Venn Diagrams 5
1.12 False positive and false discovery rate calculations 5
*2* MOCCASIN algorithm details 6
2.1 MOCCASIN Input and Output 6
2.2 Normalizing junction counts to PSI 6
2.3 Modeling and adjusting PSI using OLS regression 6
2.4 Modeling and adjustment with optional arguments 7
2.5 Computing adjusted junction counts from adjusted PSI 7
2.6 Augmentation of the model matrix to include factors from unknown confounders 8
2.7 Modeling unwanted variation using control samples 8
2.8 Modeling unwanted variation using residuals with respect to known factors 9
2.9 Discovering unknown confounding factors 9
2.10 Speed and memory optimizations 9
2.11 Majiq-specific considerations 10
3 Supplementary Figures 10
3.1 Supplementary Figure 1: Batch effect associated with different laboratories 11
3.1.1.1 Method details 11
3.1.1.2 Caption 11
3.2 Supplementary Figure 2: Batch difference FPR and tissue difference FDR with MOCCASIN discovering and removing 1, 2, or 3 unknown confounders 11
3.2.1.1 Methods 11
3.2.1.2 Caption 11
3.3 Supplementary Figure 3: Clustering of ENCODE dPSI before vs after MOCCASIN 12
3.3.1.1 Methods 12

3.3.1.2 Caption 12
4 Supplementary References 12

### 1 MOCCASIN Supplementary Methods

#### 1.1 Data Availability

The TARGET results published here are in whole or part based upon data generated by the Therapeutically Applicable Research to Generate Effective Treatments (https://ocg.cancer.gov/programs/target) initiative, phs000218. The TARGET data used for this analysis were accessed under Project #10088: Alternative splicing in pediatric cancers (request 41466-5). The SRA toolkit was used to download sra files for the TARGET dataset and simulated data was based off of data from (Zhang et al. 2014) (GEO accession GSE54652). The ENCODE project fastqs were downloaded from www.encodeproject.org (See Supplementary Table 1 for a list of file accessions used).

#### 1.2 RNA-Sequencing data processing steps

SRA files were converted to fastqs with fastq-dump (sratoolkit.2.9.2) using the following commands: --split-3 –gzip. Sequencing adaptors and low quality base calls were trimmed from fastqs using trim_galore (0.5.0) using the following commands: --stringency 5 --length 35 -q 20. For gene expression analyses, transcript per million (TPMs) and effective length quantifications were obtained using Salmon (0.11.3) in mapping-based mode with default settings. The Salmon transcriptome indices were prepared with the GRCh38 annotation for human samples and the Gencode M21 annotation for mouse samples. For alternative splicing analyses, trimmed fastqs were aligned with STAR (2.5.2a) using the following commands: --outSAMattributes All --alignSJoverhangMin 8 --readFilesCommand zcat --outSAMunmapped Within --outSAMtype BAM Unsorted. The aligned bams were sorted with samtools (1.9). The sorted bam files were then quantified for alternative splicing with MAJIQ (See 'Splicing analysis').

#### 1.3 Introducing batch effects by perturbing TPMs

Batch effects were introduced to the mouse aorta and cerebellum sample transcript level TPMs. The aorta samples used were SRR1158521, SRR1158522, SRR1158523, SRR1158524, SRR1158525, SRR1158526, SRR1158527, SRR1158528 and the cerebellum samples used were SRR1158545, SRR1158546, SRR1158547, SRR1158548, SRR1158549, SRR1158550, SRR1158551, SRR1158552. The first four of the aorta samples (SRR1158521...24) and the first four of the cerebellum samples (SRR1158545...48) were defined as "batch 1." The last four of the aorta samples (SRR1158525...28) and the last half of the cerebellum samples (SRR1158549...52) were defined as "batch 2." Batch effect perturbations were only ever introduced to the batch 2 samples and batch effects were restricted to genes with at least two protein coding transcripts and at least one transcript with > 10 reads per kilobase of transcript length in every sample. The procedure to introduce batch effects is as follows: first, the most abundant protein coding transcript per gene was identified as the transcript having the maximum over all transcripts of the minimum reads per kilobase over all samples. This definition ensures the selected transcript is not zero in any sample. Then, for a given gene a batch effect was introduced by (1) selecting a transcript uniformly at random (excluding the most abundant transcript) and (2) reducing TPM of the most abundant transcript by a factor of 'C% Change in Isoform TPM' and correspondingly increasing the TPM of the randomly selected transcript, thus maintaining the overall TPM of the gene and not breaking the definition of TPM (sum of all TPMs in a sample is one million). The 'C% Change in Isoform TPM' factors included 2%, 10%, and 60%. In addition to introducing three different levels of percent changes in isoform TPMs, we also varied the percent of genes batch-effected. We introduced batch effects to G=0%, 2%, 5% or 20% of all genes with protein coding transcripts.

#### 1.4 Simulated data generation

The mouse cerebellum and aorta simulated data are based on the real mouse cerebellum and aorta data from (Zhang et al. 2014). We used transcript-level TPM quantifications from these data as the empirical distributions for simulating RNA-Seq reads with the Benchmarker for Evaluating the Effectiveness of RNA-Seq Software (BEERS) (Grant et al. 2011). For all BEERS simulations we used the same GENCODE release M21 gene models (Frankish et al. 2019) as for transcript quantification, and the sequence from build GRCm38.p6 of the mouse reference genome (downloaded from the GENCODE website). Briefly, we directly converted the table of TPM values for each input sample into a BEERS feature quantification file so we could simulate RNA-Seq data with the same expression distributions as the input samples. Next, we used BEERS to simulate 30,000,000 paired end RNA-Seq reads for each input sample, with uniform coverage across the length of each expressed transcript and no intronic expression. BEERS introduced polymorphisms and errors into the resulting data with the following parameters: substitution frequency = 0.001, indel frequency = 0.001, error rate = 0.005. The BEERS simulator generated FASTQ files containing the simulated read pairs, a listing of the true alignments and source transcript for each read pair, and the true expression counts for each transcript in the simulated sample. To introduce systematic changes to those quantifications, we "perturbed" (see above) TPMs prior to running the BEERS algorithm. Simulated fastqs were then processed as described above.

#### 1.5 ENCODE dataset analysis

The ENCODE RNA-Sequencing data used in this paper were described in a preprint (Nostrand et al. 2018) which used 1100 ENCODE RNA-Sequencing samples generated in 61 distinct experiments (batches), whereby each batch had a control and at least one knockdown target. Every ENCODE sample, including the controls, had two biological replicates performed in every batch. 29 (548 samples) and 32 (552 samples) batches used HepG2 cells or K562 cells, respectively. For the purposes of this paper, HepG2 cells and K562 cells were considered separately when modelling and quantifying batch effects. The optimal approach to model batch effects was not obvious, so a variety of strategies were tested before we decided on an approach that is (A) easily generalizable to other studies with a similar experimental design and (B) analogous, but in a different order, to the approach taken by Van Nostrand et al. Van Nostrand et al applied the ComBat batch correction algorithm directly on sample read counts, and then averaged together the controls across batches to create two "virtual" control replicates before quantifying differential expression and differential splicing. In this paper, batch effects in all samples were modelled and removed with MOCCASIN first. After batch-correction, controls across all batches were then considered altogether in one group when quantifying differential splicing between control and knockdowns with MAJIQ see 'Splicing analysis' for more details). For differential expression analysis, control replicate 1 samples and control replicate 2 samples' gene level TPMs across all the batches were averaged together to create two "virtual" control replicates.

#### 1.6 Gene expression analysis

For gene expression level analysis of the TARGET and ENCODE datasets, transcript level TPM quantifications were collapsed into gene level TPMs with tximport (bioconductor-tximport 1.12.0). For differential expression analysis of ENCODE, DESeq2 was used to quantify the log2 fold-change (shrunk with DESeq2::fcShrink using the apelgm method) and statistical significance of differential expression between the two "virtual" control replicates and a given pair of knockdown target replicates. The DESeq2 adjusted p-value of 0.05 was the threshold for identifying a differentially expressed gene as statistically significant.

#### 1.7 Splicing analysis

MAJIQ and VOILA (v2.1-46f7268) were used to quantify splicing from sorted bam files. Based on MAJIQ's PSI quantification, two methods were subsequently used to quantify differential splicing between different experimental conditions. The first is MAJIQ's Bayesian dPSI model as described in (Vaquero-Garcia et al. 2016), which is geared to produce a set of high confidence splicing changes between groups of biological replicates. Briefly, MAJIQ dPSI quantifies the probability that a difference in splice inclusion is above some threshold with a certain confidence i.e. ($P(|dPSI| >= C1) >= C2$) for example . We used a threshold of C1 = 8%, 10%, or 20% in the results reported in the main and supplementary figures. Significant differences in splicing were those identified as having a *C2* = 95% probability of being greater than the given *C1* threshold.

The second method to detect differential splicing involved a Wilcoxon rank test combined with an optional threshold on the observed difference between the mean PSI of samples from the two groups. This approach was aimed to accommodate heterogeneous groups, as we observed in ENCODE's CTRL samples (data not shown).

#### 1.8 Uniform manifold approximation and projection (UMAP)

UMAP (McInnes, Healy, and Melville 2018) was applied to gene expression (TPM and log2 fold-change) data and splicing (E(PSI), E(dPSI), and difference in median E(PSI)) data using the R package umap (0.2.4.1) with method set to 'umap-learn', n_neighbors set to 25, and the random seed set to 924319452. UMAP projections were visualized with the R package ggplot2. See Supplementary Info 3 for specific details on how to reproduce **Fig1a-d** and **Fig3a-b**.

#### 1.9 Hierarchical clustering

For hierarchical clustering of gene expression (TPM and log2 fold-change) data and splicing (E(PSI), E(dPSI), and difference in median E(PSI)) data, the ward.D2 algorithm was applied to euclidean distances between columns and between rows. Clustering results were visualized with the R packages ComplexHeatmap and Pheatmap.

#### 1.10 Percent of total variance associated with batches

For a given numeric matrix (expression levels or PSI) the total row-wise matrix variance (V0) was quantified as the sum of squared differences from the row-wise means, divided by the total number of rows and columns. The variance due to batch (VB) or any other variable of interest was quantified as the sum of squared differences from the row-wise batch means, divided by the total number of rows and columns. The fraction of the total variance associated with the batch variable, thus, is (V0 - VB) / V0. One can apply the same principle to calculate percent of total variance due to any other factor, such as donor identity (VD), bearing in mind that VB + VD may add up to more than V0.

#### 1.11 Scatterplots and Venn Diagrams

Scatterplots with lines of best fit and pearson correlations were generated using the base R lm() method. Size-scaled Venn Diagrams were generated with R package eulerr (6.1.0).

#### 1.12 False positive and false discovery rate calculations

MAJIQ was used to quantify LSV batch differences (batch 1 vs 2) and tissue differences (aorta vs cerebellum) from the simulated data. LSVs called changing if they met thresholds (i.e. dPSI>=0.1) as specified for a given plot in this paper. The simulated data without batch effects introduced were used as ground truth to identify true positive (TP) and true negative (TN) batch and tissue differences. The following commonly used statistics were then computed:

Specificity:

$$1 - \frac{TN}{FP + TN}$$

False positive rate (FPR):

1 - Specificity

Sensitivity:

$$1 - \frac{TP}{TP + FN}$$

False negative rate (FNR):

1 - Sensitivity

False discovery rate (FDR):

$$\frac{FP}{FP + TP}$$

### 2 MOCCASIN algorithm details

#### 2.1 MOCCASIN Input and Output

MOCCASIN adjusts junction counts representing evidence for RNA splicing events in order to remove confounding variation.

The main inputs provided by the user are a model matrix and list of confounding factors (subset of the model matrix columns). The model matrix identifies the input samples to be processed by MOCCASIN. In the simplest case, each input sample is represented by a coverage table, with each row of the table representing a splice junction and the row value corresponding to read-counts providing evidence for that junction. MOCCASIN considers the fundamental splicing unit of interest to be the local splicing variation (LSV), where each LSV refers to a collection of splice junctions with either a common source or target exon (Vaquero-Garcia et al. 2016b; Norton et al. 2018). Hence, the junctions represented in the coverage table are assumed to be grouped into LSVs, with a separate data structure identifying rows with LSVs. In reality, MOCCASIN is implemented to handle the more complex case in which the coverage table includes more than one column, e.g. multiple quantifications for each splice junction corresponding to different bootstraps (please see "Majiq-specific considerations").

The main output of MOCCASIN is one set of adjusted counts corresponding to each input sample.

#### 2.2 Normalizing junction counts to PSI

MOCCASIN aggregates the junction read evidence from the input samples into a table with row dimension $M$ x $K$, where $M$ is the number of common LSV junctions and $K$ is the number of input samples.

For each LSV and sample, MOCCASIN first computes the total coverage (sum of reads over junctions). Then the counts are converted to PSI (proportion spliced-in), i.e. proportion of counts per junction; each value is in [0, 1] and the values sum over junctions to 1 for each LSV. An exception is the case in which an LSV has 0 coverage for some sample; in this case the "PSI" values are set to 0 for the junctions (and hence do not sum to 1).

#### 2.3 Modeling and adjusting PSI using OLS regression

The $M$ x $K$ junction PSI values together with the model matrix are the inputs to the adjustment step. Ordinary least squares regression is used to find estimators for the model terms. Specifically, for junctions indexed 1...$M$ and samples indexed 1...K, we model

$$PSI_{mk} = \alpha_m + X_k \cdot \beta_m + \epsilon_{mk}.$$

In the above,

- $\alpha_m$ is the intercept for junction $m$
- $X_k$ is the model matrix row corresponding to the $k$'th input sample

- $\beta_m$ are the coefficients for junction *m*
- $\epsilon_{mk}$ is the error term

OLS regression is completed independently for each junction to find estimators $\hat{\alpha}_m, \hat{\beta}_m, \widehat{\epsilon_{mk}}$ for each *k*. Thus, each OLS regression problem includes *K* observations (one per sample).

For purposes of adjusting PSI, we partition the columns of the model matrix *X* into two groups, corresponding to confounders *C* and variables-of-interest/non-confounders *V*. In the case when the known confounders are supplemented with additional factors of unwanted, the columns of *C* are partitioned into known *N* and unknown *U* confounders. Then the expression above may be rewritten

$$PSI_{mk} = \alpha_m + N_k \cdot \gamma_m + U_k \cdot \delta_m + V_k \cdot \eta_m + \epsilon_{mk}$$

Here, $\gamma, \delta, \eta$ are the coefficients corresponding to *N*, *U*, *V* respectively.

Once fit, this constitutes a learned model for junction $m$. Denote by $\hat{\alpha}_m$,$\hat{\gamma}_m$,$\hat{\delta}_m$ ,$\hat{\eta}_m$ the learned coefficients:

$$\widehat{PSI}_{mk} = \hat{\alpha}_m + N_k \cdot \hat{\gamma}_m + U_k \cdot \hat{\delta}_m + V_k \cdot \hat{\eta}_m$$

The adjustment procedure for PSI is then executed by computing:

$$r_{mk} := PSI_{mk} - \widehat{PSI}_{mk}$$
$$Z_{mk} := \hat{\alpha}_m + V_k \cdot \hat{\eta}_m$$
$$A_{mk} := r_{mk} + Z_{mk} ,$$

so that:

- $r_{mk}$ is the residual
- $Z_{mk}$ is PSI predicted from the *partial model* which excludes confounding factors
- $A_{mk}$ is the adjusted PSI

#### 2.4 Modeling and adjustment with optional arguments

The adjustment procedure above is different if the **-F (--full_correction)** argument is used. In this case, the variables of interest *V* are *not* included in the model, which results in a stronger correction, albeit one which possibly removes more variation associated with those variables.

In some cases it is desired to model variation using only control samples, which are constant with respect to the main variables of interest, and then apply the model to all samples for adjusting reads. If the user has included a **build_model** column in the model matrix, indicating a wish to model variation using only control samples, then the PSI data and model matrix are limited to those samples in the above modeling step.

#### 2.5 Computing adjusted junction counts from adjusted PSI

In the final adjustment step,

1. Negative PSI are clipped to 0
2. PSI are re-normalized for each LSV so that the sum over junctions is 1 for each sample. As before, LSVs which are all-zero (after adjustment) remain 0.

3. Adjusted counts for each junction are computed by multiplying the adjusted PSI into the initial coverage for that LSV and sample.

#### 2.6 Augmentation of the model matrix to include factors from unknown confounders

The model matrix input by the user includes filenames, variables of interest, and known confounders. The user may optionally request that MOCCASIN utilize a routine based on the RUVSeq (Risso et al. 2014) approach to incorporate additional factors of unwanted variation, by setting the -U option to 1 or more.

The routine executes one of two RUV-inspired procedures to discover factors of variation in the data which are not explained by the factors in the model matrix input by the user. It then appends the new factors to the user's model matrix and keeps the result if the matrix remains full-rank.

In order to utilize either procedure, MOCCASIN prepares a table of junction PSI values as input:

1. The junction reads for each input sample are summarized as the median over bootstrap sets for each junction, resulting in one value per junction.
2. The median junction reads are collected into a table and normalized to PSI as in the "Normalizing" section above.
3. From each LSV, the junction with the highest variance in PSI over samples is identified. The top NUM_JUNCS (default: 10,000) such junctions by variance are selected; this constitutes the input data for the following procedures.

Detail on the two procedures follows.

#### 2.7 Modeling unwanted variation using control samples

As described above in the "modeling" section, we consider three kinds of factors: main variables of interest, known confounders, and unknown confounders. In some cases it is desired to model variation using only control samples, which are constant with respect to the main variables of interest, and then apply the model to all samples for adjusting reads.

If the user has included a build_model column, indicating a wish to model variation using only control samples, then a procedure similar to RUVs is utilized: regression is performed on the control sample PSI to remove effects from the known confounding factors, and then svd is performed to discover additional factors of unwanted variation. Concretely, the procedure is:

1. Limit the input data and model matrix to the control samples (build_model column).
2. Perform OLS regression on the data with this model matrix.
3. Compute the residuals from this regression.
4. Mean-center the residuals (from each value, subtract the junction mean over samples).
5. Compute the singular value decomposition $USV^T=Y_{rc}$, where $Y_{rc}$ is the matrix of centered residuals.
6. Compute the factor loadings: $A := SV^T$
7. Estimate the factors of unwanted variation by ordinary least squares regression of the factor loadings on all samples: $W := Y^T A^T (AA^T)^{-1}$, where Y denotes the PSI matrix with all samples included.

8. Append the estimated factors (up to the number specified by the -U argument) to the original model matrix and retain them if the model remains full-rank.

#### 2.8 Modeling unwanted variation using residuals with respect to known factors

Otherwise, a procedure similar to RUVr is utilized: regression is performed on all the samples to remove effects from the known confounding factors and main variables of interest, and then svd is performed to discover additional factors. All factors supplied by the user (confounders and non-confounders) are included in the model to ensure that new factors discovered by this procedure correspond to variation not already explained by known factors. Concretely, the procedure is:

1. Perform OLS regression on the input data Y with this model matrix.
2. Compute the residuals $Y_r$ from this regression.
3. Compute the singular value decomposition: $WSV^T=Y_r$
4. Append the estimated factors W (up to the number specified by the -U argument) to the original model matrix and retain them if the model remains full-rank.

#### 2.9 Discovering unknown confounding factors

Our method for discovering unknown confounding factors is based on that by Risso et al with some modifications. We implemented these routines because we need to initially model and take out the effects of the *known* confounding factors so that these are not rediscovered by the RUVs procedure. (The RUVs procedure assumes that only two kinds of effects exist, termed "main variables of interest" and "unwanted variation"; it does not handle known confounders, and as currently implemented would rediscover known confounders corresponding to a large proportion of the variation in the data.) Hence our inputs to RUVs are residuals, which may be negative; since RUVs expects counts and tries to take their logarithm, we found RUVs unworkable for our purposes. Therefore, our RUVs-style procedure includes an initial step which computes the residuals of an OLS regression on the input PSI with respect to known confounding factors. Our procedure also works on PSI rather than log counts, although it could execute just as well with either.

In summary, our RUVr-like procedure is based on RUVSeq except that the step to generate the residuals utilizes OLS regression instead of a negative binomial GLM suggested by the RUV documentation (we use PSI rather than counts). We also note that, like the RUVr procedure as implemented currently in RUVSeq (July 2019), our derived factors correspond to only the left singular vectors in the singular value decomposition; this implementation contrasts with the method reported in the 2014 Nat Biotech paper, which states that the derived factors should be the left singular vectors composed with the singular values.

Both routines are implemented in python as part of the MOCCASIN module.

#### 2.10 Speed and memory optimizations

In contrast to gene or transcript expression studies, which typically utilize tens or at most a few hundred thousand tags, MOCCASIN must model and adjust counts for possibly millions of splice junctions in a single run. The size of the data necessitates additional speed and memory optimizations.

MOCCASIN performs linear regression on PSI. While this may seem like a simple first-pass idea, we note that the model was originally for log counts as a fast approximation to inference with e.g. a negative binomial GLM used for count data in applications such as DESeq2 (Love et al, Genome Biol. 2014).

MOCCASIN partitions LSVs into "chunks" for processing in order to limit peak memory usage. This is possible because LSVs are modeled and adjusted independently of one another in the MOCCASIN framework. The user can set the max chunk size in terms of the max number of coverage table cells to be loaded at once into memory (-M). However, the peak memory use currently is lower-bounded by the size of the coverage table in the largest input .majiq file.

MOCCASIN allows the user to specify the number of processes to use (-J) as a speed optimization. This governs the number of spawned processes which will process each chunk of LSVs in parallel during the modeling and adjustment steps.

For large (i.e. >500 samples) datasets, I/O becomes a rate-limiting process since we try to keep memory use low (see above) by writing and reading temporary files to a user-specified tmp location. The temporary files would not be necessary if we made use of shared memory across processes. Python3.8 does include shared memory capabilities but python3.8 is not yet as widely used as python3.6 and python 3.6 lacks shared memory capabilities. To facilitate MOCCASIN usability, we chose to use python3.6, and to make up for lack of shared memory in python3.6, we suggest users point the MOCCASIN tmp directory to the fastest disk available (e.g. a RAM drive at /dev/shm). At the end of the MOCCASIN execution, the tmp subdirectory is automatically deleted (unless the -K flag is supplied).

#### 2.11 Majiq-specific considerations

The previous sections described a simple case in which each input sample includes one value (e.g. "read counts") per splice junction. MOCCASIN is written to work with the output of a MAJIQ build (Vaquero-Garcia J. et al, ELife 2016;10.7554/eLife.11752, Norton S. et al 2017). In reality, MAJIQ outputs many (default: 30) values per splice junction. These values are *read rates*, which are the estimated number of junction reads per position for nucleotide positions near the splice junction; MAJIQ bootstraps over junction-proximal nucleotide positions and includes read rates corresponding to many bootstrap samples. MOCCASIN considers each bootstrap sample independently and thus models and adjusts read rates (rather than counts) for each bootstrap sample separately.

### 3 Supplementary Figures

The subsections below include the caption for each of the supplementary figures (separate files) along with the specific methods used to derive those.

#### 3.1 Supplementary Figure 1: Batch effect associated with different laboratories

###### 3.1.1.1 Method details

RNA-Seq of mouse hippocampus from (Peixoto et al. 2015) were downloaded with fastq-dump v2.9.2 from GEO (Wood lab, GSE44229; Abel lab, GSE63412). Reads were trimmed with trimgalore v0.4.4_dev, aligned to Gencode M20 using STAR v2.6.1a, and analyzed for splicing with MAJIQ v2. Samples SRRs processed included: SRR7072**19-24**, 6 samples, Wood lab Object-Location Memory (OLM) treatment; SRR7072**25-30**, 6 samples, Wood lab, control treatment, SRR16566**67-71**, 5 samples, Abel lab, control treatment; SRR16566**73-76**, 4 samples, Abel lab, Fear Conditioning (FC) treatment. MOCCASIN model matrix included: intercept, Treatment_OLM, Treatment_FC. No explicit confounder factor was included even though we know the Lab_Wood / Lab_Abel difference. Instead, we used MOCCASIN -U (see methods above) with 2 hidden factors. For evaluation, 10,000 LSVs were selected uniformly at random from those originally quantified in all samples. Principal component analysis (PCA) from scikit-learn was applied to PSI from the 10,000 LSVs, and plots were generated with matplotlib. MAJIQ was used to quantify differences in splicing between Wood vs Abel control samples and WILCOXON test applied to quantify p-values.

###### 3.1.1.2 Caption

**MOCCASIN removes batch effects associated with different laboratories**. PCA of PSI shows samples cluster by batch before MOCCASIN in **a**, then by treatment after MOCCASIN as shown in **b**. **c**, quantification of false positive differences between Wood vs Abel laboratory's control samples before (blue) vs after (orange) MOCCASIN. **d**, PSI of control samples before (original) vs after (adjusted) MOCCASIN. OLM, Object-oriented learning; FC, Fear Conditioning.

#### 3.2 Supplementary Figure 2: Batch difference FPR and tissue difference FDR with MOCCASIN discovering and removing 1, 2, or 3 unknown confounders

###### 3.2.1.1 Methods

Used MAJIQ dPSI to quantify batch differences (batch 1 vs 2) and tissue differences (aorta vs cerebellum) from the simulated data. LSVs called changing if they met the threshold Probability(|E(dPSI)|>=0.2)>=0.95. See 1.12 on how false positive rate (FPR) and false discovery rate (FDR) were calculated.

###### 3.2.1.2 Caption

Batch 1 vs batch 2 false positive rate (FPR) or aorta vs cerebellum false discovery rate (FDR) from the batch-effected simulated data (blue, before MOCCASIN) versus the corrected data (orange or grey, after MOCCASIN). MOCCASIN was run in three different configurations: 1 known confounding factor provided to MOCCASIN (orange), or 1, 2, or 3 (gradient of light-to-dark greys) unknown confounding factors identified and removed by MOCCASIN.

#### 3.3 Supplementary Figure 3: Clustering of ENCODE dPSI before vs after MOCCASIN

###### 3.3.1.1 Methods

MAJIQ was used to quantify changes in splicing (dPSI) between controls and target knockdowns (See 1.5 above). Including HepG2 and K562, there were 1100 control vs target comparisons. First, MAJIQ results from all 1100 comparisons were combined into one table (Supplementary Info 2), next these junctions were filtered: at least one comparison had a |dPSI|>=0.1 and TNOM P-value less than 0.05, then for each LSV, the junction with the greatest |dPSI| value kept, breaking ties randomly. R package pheatmap v1.0.12 was used to generate the clustergrams (distance metric: 'euclidean', clustering method: 'complete'.)

###### 3.3.1.2 Caption

Each row is an ENCODE comparisons' dPSI values. The clustergrams on the left indicate euclidean distances between comparisons. Each colored bar provides information about the given comparison: the sequencing platform (HiSeq 2000, black; HiSeq 2500, grey), the batch identifier (1:61, rainbow-colored), the type of experiment (shRNA, black; CRISPR knockout, red), and the cell type (HepG2, yellow; K562, pink). The left figure is before MOCCASIN, and right is after MOCCASIN. Note that after MOCCASIN, Batch IDs colors no longer segregate together, rather, the colors are diapered randomly.
