## Supplementary figures and images for "MOCCASIN: A method for correcting for known and unknown confounders in RNA splicing analysis"

### Supplemental Figures

#### Supplemental Figure 1

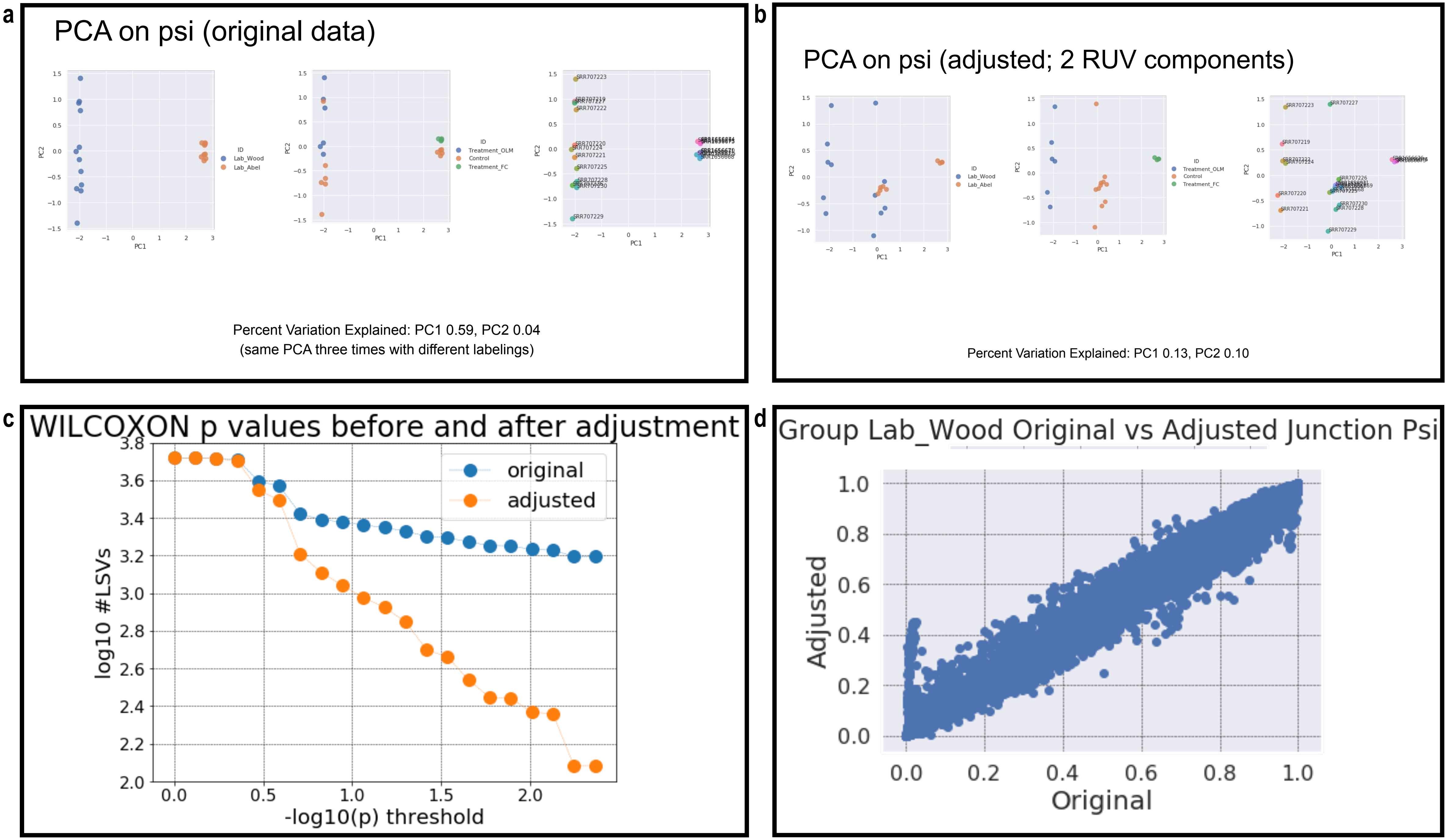

#### Supplemental Figure 2

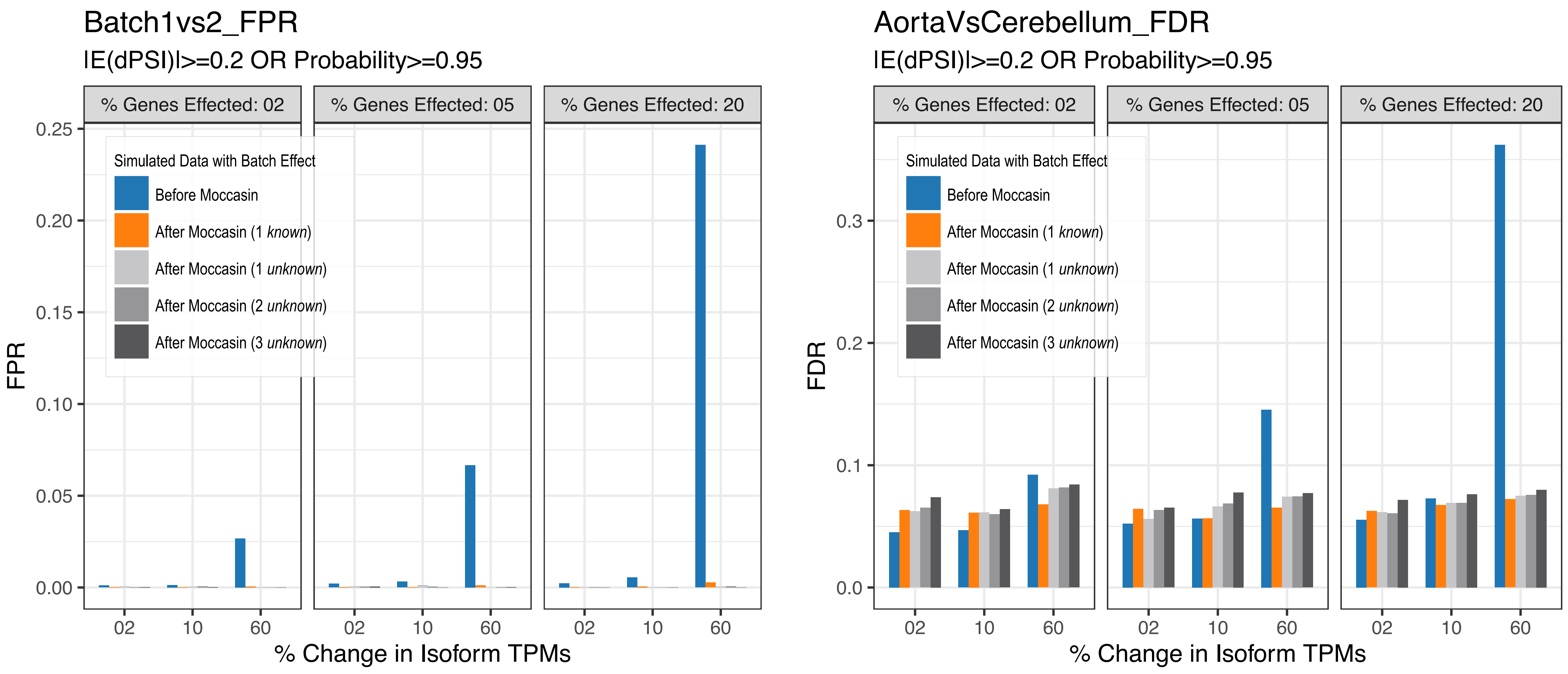

**Supplemental Figure 3**

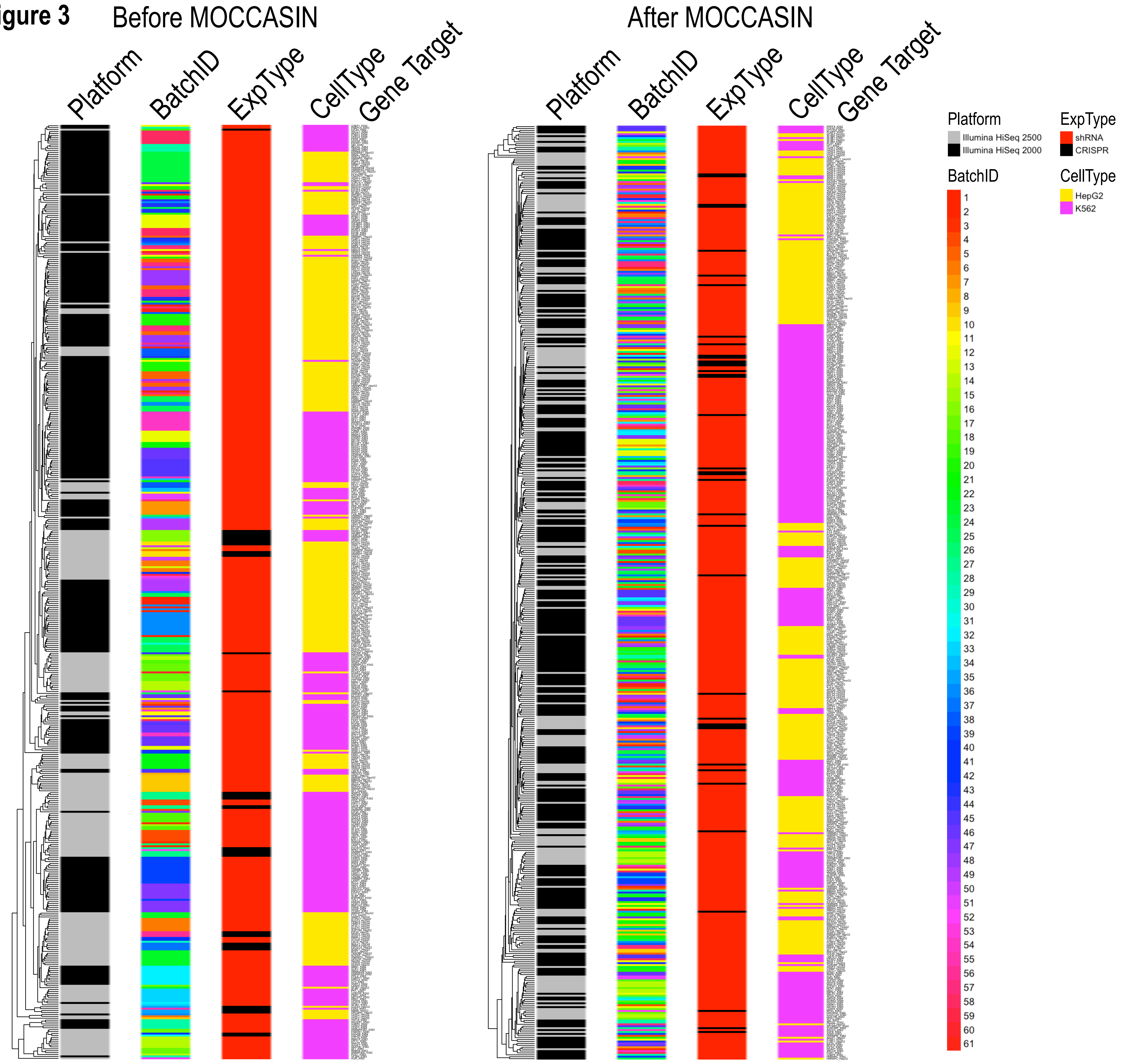
